# Cefiderocol is an effective topical monotherapy for experimental extensively-drug resistant *Pseudomonas aeruginosa* keratitis

**DOI:** 10.1101/2023.08.31.555778

**Authors:** Eric G. Romanowski, Sonya M. Mumper, Hazel Q. Shanks, Kathleen A. Yates, Jonathan B. Mandell, Michael E. Zegans, Robert M. Q. Shanks

## Abstract

**Purpose:** To test cefiderocol, a siderophore-cephalosporin antibiotic for topical monotherapy treatment of experimental extensively drug resistant (XDR) *Pseudomonas aeruginosa* keratitis.

**Design:** Preclinical study.

**Subjects and Controls:** Deidentified *P. aeruginosa* keratitis isolates, XDR *P. aeruginosa* from eye drop outbreak, rabbits, saline, cefiderocol 50 mg/ml, ciprofloxacin 0.3%, and tobramycin 14 mg/ml.

**Methods, Intervention, or Testing:** Cefiderocol antibacterial activity against *P. aeruginosa* keratitis isolates (n=135) was evaluated by minimum inhibitory concentration (MIC) testing. Ocular toxicity/tolerability and antibacterial efficacy were tested *in vivo* with experimental rabbit models. Corneal concentrations and stability were assessed using a bioassay.

**Main Outcome Measures:** MIC analysis for susceptibility, graded tests for ocular toxicity/tolerability, CFU analysis for bacterial burden, corneal cefiderocol concentrations.

**Results:** 100% of *P. aeruginosa* keratitis isolates were susceptible to cefiderocol (n=135), the MIC_90_ was 0.125 µg/ml including the XDR isolate (MIC = 0.125 µg/ml). Topical cefiderocol 50 mg/ml was minimally toxic to the ocular surface and was well tolerated. For the XDR *P. aeruginosa* isolate, topical cefiderocol 50 mg/ml, significantly decreased corneal CFU compared to ciprofloxacin 0.3%, tobramycin 14 mg/ml, and saline. In addition, tobramycin 14 mg/ml was more effective than the saline control. Mean cefiderocol corneal concentrations were 191x greater than the MIC_90_ of the *P. aeruginosa* keratitis isolates. Refrigerated cefiderocol maintained antimicrobial activity over a one-month period.

**Conclusions:** These results demonstrate that cefiderocol is well tolerated on rabbit corneas and is effective against *P. aeruginosa* keratitis isolates *in vitro* and was effective *in vivo* against an XDR isolate in a rabbit keratitis model. Given the recent outbreak of keratitis caused by this XDR *P. aeruginosa*, cefiderocol is a promising additional antibiotic that should be further evaluated for topical treatment of keratitis caused by antibiotic resistant *P. aeruginosa*.

## INTRODUCTION

The Center for Disease Control and Prevention (CDC) and numerous news organizations recently reported a dangerous outbreak of eye infections linked to the use of artificial tears purchased online ^1-4^. Patients identified in this outbreak have experienced serious corneal infections which have led to blindness, loss of eyes, and even death in several patients. Notably, the *Pseudomonas aeruginosa* bacteria causing the infections was extensively drug resistant (XDR) including to all antibiotics currently used to topically treat eye infections ^2, 3^. This strain has the Verona integron-mediated metallo-β-lactamase and Guiana extended-spectrum β-lactamase genes ^1, 4, 5^.

The World Health Organization and CDC consider antibiotic resistant bacteria to be a global threat ^6^. Keratitis patients infected with multidrug resistant bacteria have worse clinical outcomes and longer and more costly treatment ^7-9^. While XDR keratitis isolates are unusual, the antibiotic resistance in ophthalmology monitoring study, ARMOR, has documented a concerning frequency of antibiotic resistant keratitis isolates ^10, 11^.

Because of the existing threat of XDR pathogens, there is a need to identify new antimicrobials for the topical treatment of keratitis. The latest antibiotic FDA approved for treatment of eye infections, besifloxacin, was developed in the 1990s. The major antibiotics (aminoglycosides and fluoroquinolones) used for treatment of bacterial keratitis were approved in the 1970s through the early 2000s. Fluoroquinolone and aminoglycoside class antibiotics have concerning side effects and increasing rates of bacterial resistance leading to reduced use for systemic infections ^12-14^. The more recent ocular antibiotics, such as moxifloxacin and besifloxacin, were designed for better efficacy against Gram-positive bacteria. As such, there is a clear need for newer approaches to treat keratitis caused by Gram-negative bacteria.

Cefiderocol is a rationally designed antibiotic that combines a cephalosporin antibiotic and siderophore-like iron binding moiety into one molecule (Figure 1 Chemical Structure) ^15, 16^. It effectively tricks bacteria into taking up the antibiotic due to the siderophore component, so it has been called a Trojan horse strategy antibiotic ^17^. The commercial formulation of cefiderocol, FETROJA, was US FDA approved in 2019 for the treatment of complicated urinary tract infections caused by Gram-negative bacteria ^18^. Specifically, urinary tract infections caused *Enterobacter cloacae* complex, *Escherichia coli*, *Klebsiella pneumoniae*, *Proteus mirabilis*, and *Pseudomonas aeruginosa*, and nosocomial pneumonia caused *by Acinetobacter baumannii*, *Enterobacter cloacae* complex, *Escherichia coli*, *Klebsiella pneumoniae*, *Pseudomonas aeruginosa*, and *Serratia marcescens*. Cefiderocol was the only antibiotic to which artificial tear associated outbreak XDR *P. aeruginosa* bacteria was susceptible *in vitro* ^1^; however, it has not been tested for topical use in the eye.

**Figure 1.**
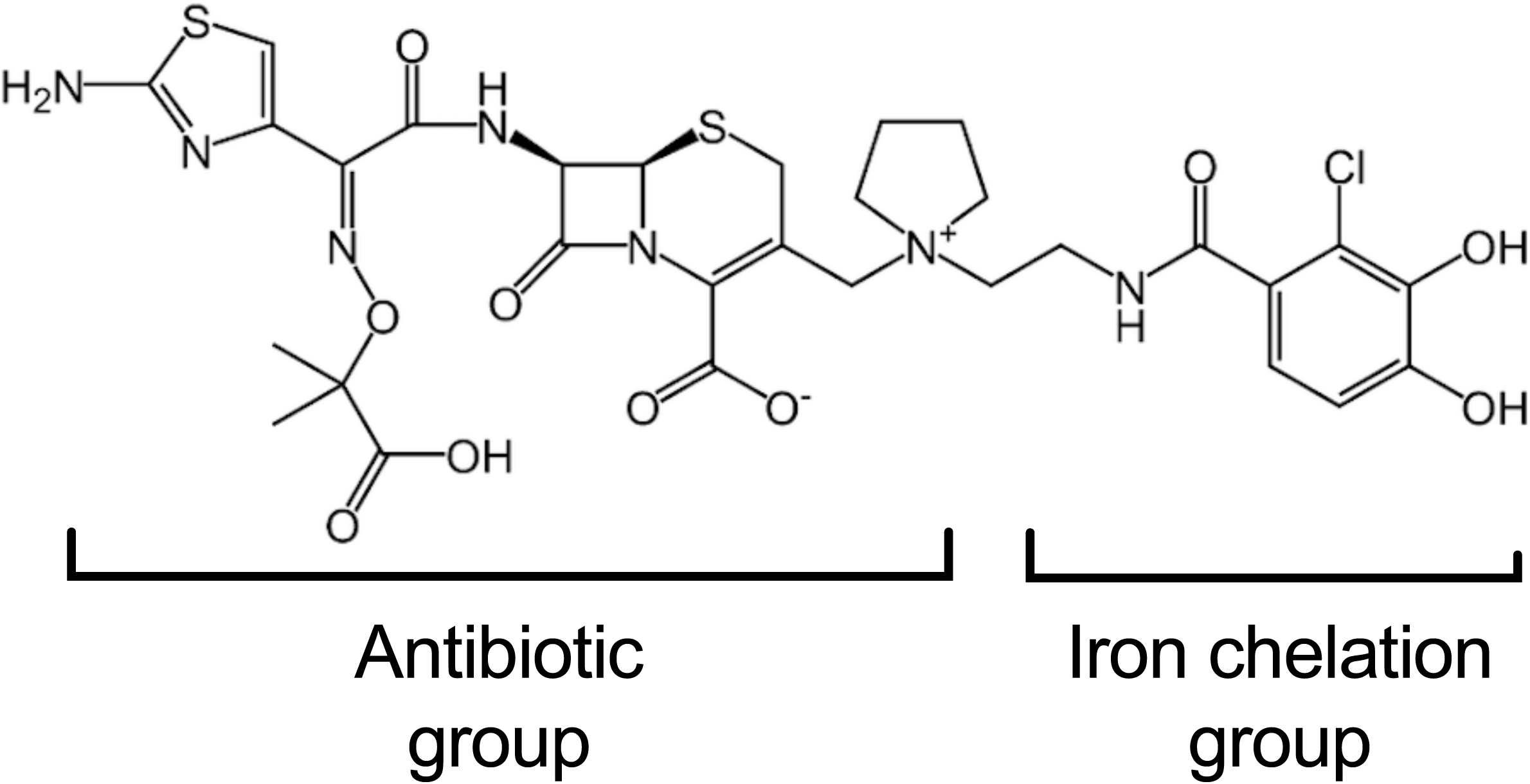
Structure of cefidericol.

The goal of this study is to determine the suitability of cefiderocol as a topical antibiotic for the treatment of experimental *P. aeruginosa* keratitis. Toward this objective, the susceptibility of a panel of keratitis isolates to cefiderocol was evaluated *in vitro*. The tolerability of several concentrations of cefiderocol was determined after topical application to New Zealand White (NZW) rabbit eyes. Finally, the ability of highest tolerable concentration of topical cefiderocol to treat *P. aeruginosa* keratitis caused by an antibiotic susceptible keratitis isolate and the XDR strain from the artificial tear outbreak (obtained from the CDC) were evaluated in an experimental NZW rabbit keratitis model. The efficacy of cefiderocol was compared to standard of care antibiotics ciprofloxacin 0.3% and fortified tobramycin 14 mg/ml. The stability of the topical cefiderocol solutions was evaluated over the course of a month. Lastly, the cefiderocol concentrations in rabbit corneas was evaluated using a bioassay. Together the data from this study support the idea that cefiderocol should be further evaluated for use in human patients for *P. aeruginosa* keratitis caused by an XDR strain.

## METHODS

### Bacterial Strains

For *in vitro* Minimum inhibitory concentration (MIC) determinations, deidentified *P. aeruginosa* strains were isolated from patients with keratitis presenting to the Charles T. Campbell Ophthalmic Microbiology Laboratory at the Department of Ophthalmology at the University of Pittsburgh School of Medicine. For the *in vivo* studies, two strains of *P. aeruginosa* were used. Strain K900 is a keratitis isolate previously used in experimental keratitis studies ^19-21^, and is a cytotoxic ExoU+/ExoS- strain based on its genome sequence (Genbank NZ_JAODAA000000000). Strain CDC1270 is an XDR isolate that was obtained from the CDC. Strain CDC1270 was isolated from the cornea of a patient from the recent artificial tears eye infection outbreak ^1, 4, 5^. Isolates of the outbreak strain have been sequenced revealing multiple resistance determinants ^2, 5^.

### Antibiotic Susceptibility Testing

MIC testing of the deidentified *P. aeruginosa* keratitis isolates for cefiderocol and other standard of care antibiotics used to treat *P. aeruginosa* keratitis (tobramycin, ciprofloxacin, and ceftazidime) was performed using MIC strips (LIOFILCHEM, Waltham, MA) as previously described ^22^. Resistance was determined using Clinical and Laboratory Standards Institute (CLSI) systemic breakpoints ^23^. The MIC values for the strains used in the *in vivo* studies, K900 and CDC1270, are presented in Table 1.

**Table 1.** Susceptibility of *P. aeruginosa* strains used in this study.

| Antibiotic | K900<br>(MIC µg/ml) | CDC1270<br>(MIC µg/ml) |
| --- | --- | --- |
| Cefiderocol | 0.023 (S) | 0.125 (S) |
| Ceftazidime | ND | >256 (R) |
| Ciprofloxacin | 0.19 (S) | >32 (R) |
| Tobramycin | 0.5 (S) | >256 (R) |

ND, not determined; (S), Susceptible; (R), Resistant

### Experimental Drugs

All drugs were purchased from the UPMC Presbyterian inpatient pharmacy. The saline used in this study was 0.9% Sodium Chloride Injection USP (saline) (Baxter Healthcare Corp. Deerfield, IL). To prepare cefiderocol, one gram vials of the parenteral version of cefiderocol, FETROJA (Cefiderocol for Injection, Shionogi Inc. Florham Park, NJ) were used. The vials were stored at 4^°^ C as directed until reconstituted. A fresh preparation of cefiderocol was made for each *in vivo* experiment. On the morning of each experiment, a 1 g vial of FETROJA was reconstituted with 10 ml of saline, and the vial was gently shaken to dissolve as per the manufacturer’s instructions. The vial was allowed to stand until the foaming generated on the surface had disappeared (typically within 2 minutes). The reconstituted solution had a final volume of approximately 11.2 ml and a concentration of 89 mg/ml. 8.43 ml of the 89 mg/ml cefiderocol stock was added to 6.57 ml of sterile saline to yield 15 ml of 50 mg/ml cefiderocol. The pH of this concentration was measured at 5.36 (Corning pH/ion analyzer 350, Corning, Corning NY). For the tolerability study, subsequent dilutions of the 50 mg/ml cefiderocol formulation were made in 0.9% sodium chloride injection, USP to produce concentrations of 25 mg/ml, 10 mg/ml, and 5 mg/ml. For all *in vivo* experiments, the cefiderocol solutions were kept on ice and out of light in a foil covered tube at all times. Following the experiments, the remaining cefiderocol solutions and any aliquots were refrigerated and were used in subsequent stability testing experiments.

For each keratitis trial, tobramycin 14 mg/ml was prepared from a 40 mg/ml stock of TOBRAMYCIN INJECTION USP (Fresenius Kabi USA, LLC, Lake Zurich IL). 3.5 ml of 40 mg/ml tobramycin was added to 6.5 ml of saline to yield 10 ml of 14 mg/ml of tobramycin. The solution was kept on ice during the experiment. 5 ml bottles of Ciprofloxacin Hydrochloride Ophthalmic Solution, 0.3% (Sandoz Inc. Princeton, NJ) were used. The ciprofloxacin solution contains 0.006% benzalkonium chloride as a preservative. The ciprofloxacin bottles were kept at room temperature as directed. Saline served as the negative control and was kept on ice during dosing.

The cefiderocol, tobramycin, and saline solutions were instilled using a Rainin E4 XLS electronic pipet set in the multi-dispense mode. 37 μl drops were instilled. Ciprofloxacin 0.3% was instilled using its commercial dropper bottle.

### Animals

NZW female SPF rabbits weighing 1.1-1.4 kg were obtained from Charles River Laboratories Canadian rabbitry. These studies conformed to the ARVO Statement on the Use of Animals in Ophthalmic and Vision Research and were approved by the University of Pittsburgh’s Institutional Animal Care and Use Committee (IACUC Protocol 23053154).

### Ocular Toxicity/Tolerability Study

An ocular toxicity/tolerability study was performed to determine whether several concentrations of topical cefiderocol were toxic to the ocular surface and whether the topical drops were tolerable. Fifteen NZW rabbits underwent a slit-lamp examination on the day before the ocular toxicity/tolerability study to determine whether there were any preexisting abnormalities. On the day of the study, the 15 rabbits were divided into 5 groups of 3 rabbits each: 1) Cefiderocol 50 mg/ml; 2) Cefiderocol 25 mg/ml; 3) Cefiderocol 10 mg/ml; 4) Cefiderocol 5 mg/ml; 5) Saline. Topical 37 μl drops were instilled into both eyes every 30 minutes for 6 hours (13 total doses). During each instillation, the behavior of the rabbits was documented and given a numerical score based on their behavior. The scoring system used was the following: no reaction after instillation (0), closed eyes after instillation (1); demonstrated delayed eye wiping (10 – 30 seconds) after instillation (2); demonstrated immediate eye wiping after instillation (3); flinching after instillation (4); vocalizing after instillation (5). Following the final dose, the eyes were examined using the slit-lamp and graded using the Modified MacDonald-Shaddock Scoring System ^24^. The eyes were again examined and graded 3 days later to determine whether any delayed toxicity was produced.

### *P. aeruginosa* NZW Rabbit Keratitis Model

A bacterial keratitis model optimized for evaluation of antimicrobials was used as previously described ^25, 26^. A total of 30 NZW rabbits were used in duplicate trials of 15 rabbits for experiments involving each of the two *P. aeruginosa* strains, K900 and the CDC1270. The 30 total rabbits for each *P. aeruginosa* strain were divided into 5 groups of 6 rabbits.

The rabbits were anesthetized with 40 mg/kg of ketamine and 4 mg/kg of xylazine administered intramuscularly. The corneas of the left eyes were anesthetized with topical 0.5% proparacaine. The eyes were proptosed and 6 mm areas of the central corneal epithelium of the left eyes were removed using an Amoils epithelial scrubber (Innovative Excimer Solutions, Toronto, ON, Canada). The creation of the epithelial defect was to mimic the large epithelial defects demonstrated in patients with XDR *P. aeruginosa* keratitis ^2^. Immediately afterward, the corneas were injected intrastromally with the 25 µl of phosphate buffered saline (PBS) containing *P. aeruginosa* (∼2000 colony forming units [CFU]) below the area from which the epithelium was removed. Rabbits were treated with intramuscular doses of 1.5 mg/kg of ketoprofen for analgesia.

Actual inocula for each trial was determined using the EddyJet 2 spiral plating system (Neutec Group Inc., Farmingdale, NY) on 5% trypticase soy agar with 5% sheep’s blood plates. Plates were incubated for ∼18-20 h at 37°C and the colonies were enumerated using the Flash and Grow colony counting system (Neutec Group, Inc).

Sixteen hours after inoculation, the rabbits were divided into 5 groups (n=6): 1) Cefiderocol 50 mg/ml; 2) Ciprofloxacin 0.3% (Standard of Care Control); 3) Tobramycin 14 mg/ml (Standard of Care Control); 4) Saline (Negative Control); and 5) No Treatment (Onset of Therapy Control). Topical therapy for all groups was initiated at this time. The treatment regimen consisted of 1 drop in both eyes every 15 minutes for 1 hour, then every 30 minutes for 7 hours (19 total doses over 8 hours).

The final group (Group 5) was euthanized at 16 hours post-inoculation and the corneas were harvested prior to drug treatment to establish baseline colony count determinations at the onset of therapy. The rabbits were systemically anesthetized with ketamine and xylazine as described above and euthanized with an intravenous overdose of Euthasol solution (390 mg/ml pentobarbital sodium, 50 mg/ml phenytoin sodium) following the 2020 AVMA Euthanasia Guidelines. Corneal buttons were harvested from the area encompassing visible corneal opacities using a 10 mm trephine and placed into Lysing Matrix A tubes (MP Biomedicals) containing 1 ml of PBS. The corneas were then homogenized with an MP Fast Prep-24 homogenizer (MP Biomedicals), and the numbers of corneal bacteria were enumerated as described above.

Prior to the final topical treatments, the rabbits’ eyes were examined using a slit-lamp and photographed using the camera on the slit-lamp. The Onset of Therapy groups were also similarly examined and photographed prior to euthanasia. Following the final topical treatment, the rabbits were euthanized, the corneas harvested and homogenized, and colony count determinations performed as described above. The areas of the corneal infiltrates produced for each treatment group and *P. aeruginosa* strain were determined using ImageJ software (NIH) from photographs taken of the eyes at a consistent distance and magnification.

### Semi-quantitative Bioassay for Cefiderocol Stability and Corneal Concentrations

The stability of the topical 50 mg/ml concentrations of cefiderocol was determined using a bioassay. After each rabbit experiment (ocular toxicity/tolerability and 4 bacterial keratitis treatment trials), aliquots of the 50 mg/ml cefiderocol solutions were refrigerated. On the day of the final bacterial keratitis study, a bioassay was performed in duplicate comparing the sizes of zones of inhibition produced from previously prepared solutions to the freshly prepared solution.

On the day of the assay, 50 mg/ml cefiderocol solutions prepared 33, 27, 21, and 7 days earlier and a freshly prepared 50 mg/ml solution were serially diluted in sterile saline to produce cefiderocol concentrations of 5, 0.5, 0.05, and 0.005 mg/ml. *P. aeruginosa* strain K900 was grown overnight at 37°C on 5% trypticase soy agar with 5% sheep’s blood plates. Several colonies of *P. aeruginosa* strain were picked up with a sterile Dacron swab and 10 ml of sterile saline was inoculated with the bacteria until it was turbid. A new sterile Dacron swab was used to inoculate Mueller-Hinton agar plates to create lawns of bacteria over the entire surface of the plates. Four blank 6 mm filter paper disks were placed onto the surface of each plate. The 4 disks per plate were then inoculated with 20 µl of each of the test cefiderocol concentrations of 5, 0.5, 0.05, and 0.005 mg/ml and the disks were allowed to dry. The plates were incubated overnight at 37°C. The following day, the zones of inhibition for each concentration and preparation were measured in mm and compared. The assay was performed in duplicate for each preparation.

To determine approximate cefiderocol corneal concentrations, corneas from both the K900 and CDC1270 keratitis studies (n=12) were used in this analysis as they were all treated with the same regimen of 50 mg/ml cefiderocol. Following homogenizing and processing of the corneal buttons for colony count determinations, the supernatants of the homogenates of the cefiderocol treated corneas were processed to determine approximate cefiderocol concentrations in the homogenates. The homogenates were centrifuged and filtered through Costar Sin-X, 0.22µm cellulose acetate filters, to remove the live bacteria. Samples were either assayed at that time or were refrigerated at 4°C prior to testing for antibiotic activity. For this assay, a 50 mg/ml (50,000 µg/ml) cefiderocol solution was diluted in sterile saline to final concentrations of 50, 25, 12.5, 6.25, 3.125, and 1.56 µg/ml. These concentrations served as the standard concentrations for the assay. Mueller-Hinton agar plates were prepared as in the Cefiderocol Stability Bioassay above. Blank disks were inoculated with 20 µl of each of the standard cefiderocol concentrations and the undiluted corneal homogenate filtrates. The plates were incubated, and the zones of inhibition measured as described above. The zone sizes were modified to subtract the 6 mm disk size. The modified zone sizes of the corneal homogenate solutions were inserted into concentration equations produced using a Linear Fitted Line Plot Regression Analysis of cefiderocol concentration vs. zone size for the cefiderocol standards (Minitab Version 19, State College, PA) to determine the approximate active cefiderocol concentrations contained in the corneal homogenates.

### Statistical Analysis

The MIC data was placed into a GraphPad Prism worksheet and the median, mode, range, MIC_50_ and MIC_90_ determinations were calculated. The modified MacDonald-Shaddock scores and the behavioral scores from the Ocular Toxicity/Tolerability study were analyzed non-parametrically using Kruskal-Wallis ANOVA with Dunn’s multiple comparisons test (GraphPad Prism). The corneal colony counts + 1 were log_10_ transformed and analyzed using ANOVA with Tukey’s multiple comparisons test (GraphPad Prism) for test agent analysis.

## RESULTS

### Antibiotic Susceptibility Analysis of *P. aeruginosa* Keratitis Isolates to Cefiderocol and Standard of Care Antibiotics

The MIC values of keratitis isolates to cefiderocol were evaluated and all were found to be susceptible (Table 2 and Figure 2). There was a significantly higher frequency of isolates susceptible to cefiderocol than the other tested antibiotics including *P. aeruginosa* keratitis standard of care antibiotics ciprofloxacin (93.3% susceptible) and tobramycin (28%). The cephalosporin ceftazidime that has been suggested as an alternative to ciprofloxacin and tobramycin for treating keratitis caused by resistant isolates also had a significantly lower frequency of susceptibility compared to cefiderocol (94.9%).

**Figure 2.**
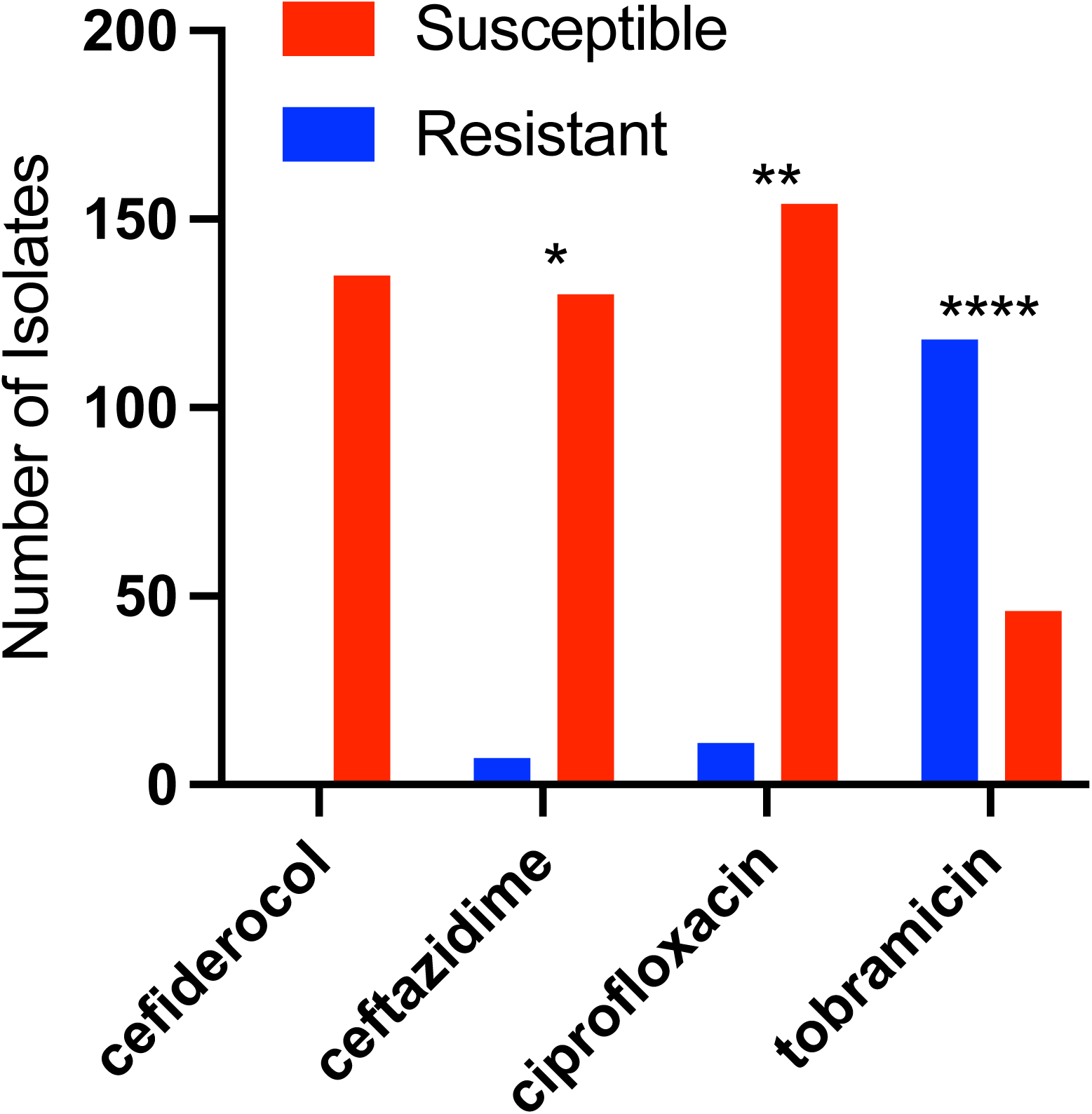
Susceptibility of *P. aeruginosa* to selected antibiotics based on MIC values compared to CLSI break points. Asterisks indicated significant differences from cefiderocol group by chi-square with Yates correction. *, p<0.05; **, p<0.01; ****, p<0.0001.

**Table 2.** Susceptibility of *P. aeruginosa* keratitis isolates. MIC values are expressed in µg/ml.

| Antibiotic | n= | Median | Mode | MIC<sub>50</sub> | MIC<sub>90</sub> | Range | % Susceptible |
| --- | --- | --- | --- | --- | --- | --- | --- |
| Cefiderocol | 135 | 0.094 | 0.064 | 0.094 | 0.125 | <0.016-0.94 | 100% |
| Ceftazidime | 137 | 2.0 | 1.5 | 2.0 | 8.0 | 0.75->256 | 94.9% |
| Ciprofloxacin | 164 | 0.19 | 0.19 | 0.19 | 0.5 | 0.047-1.5 | 93.3% |
| Tobramycin | 164 | 1.5 | 1.5 | 1.5 | 3.0 | 0.047-8 | 28% |

Susceptibility was determined using CLSI breakpoints<sup>23</sup>.

### Ocular Toxicity and Tolerability of Topical Cefiderocol

The effect of four different concentrations of cefiderocol for ocular toxicity was measured using the Modified MacDonald Shadduck scoring system. The results are divided into conjunctival scores with a 10 point total possible score and corneal scores with a 16 point possible score. Following instillation into both eyes every 30 minutes for 6 hours (13 total doses). There was a dose dependent increase in scores within the cefiderocol treatment groups for the conjunctival scores (congestion, chemosis, and discharge) but the differences were not significant (Figure 3A). As seen in Figure 3B, there were no clinical signs of corneal pathology seen and there was no fluorescein staining demonstrated in any of the corneas. Photographs of all eyes on the day of treatment are presented in Figure S1.

**Figure 3.**
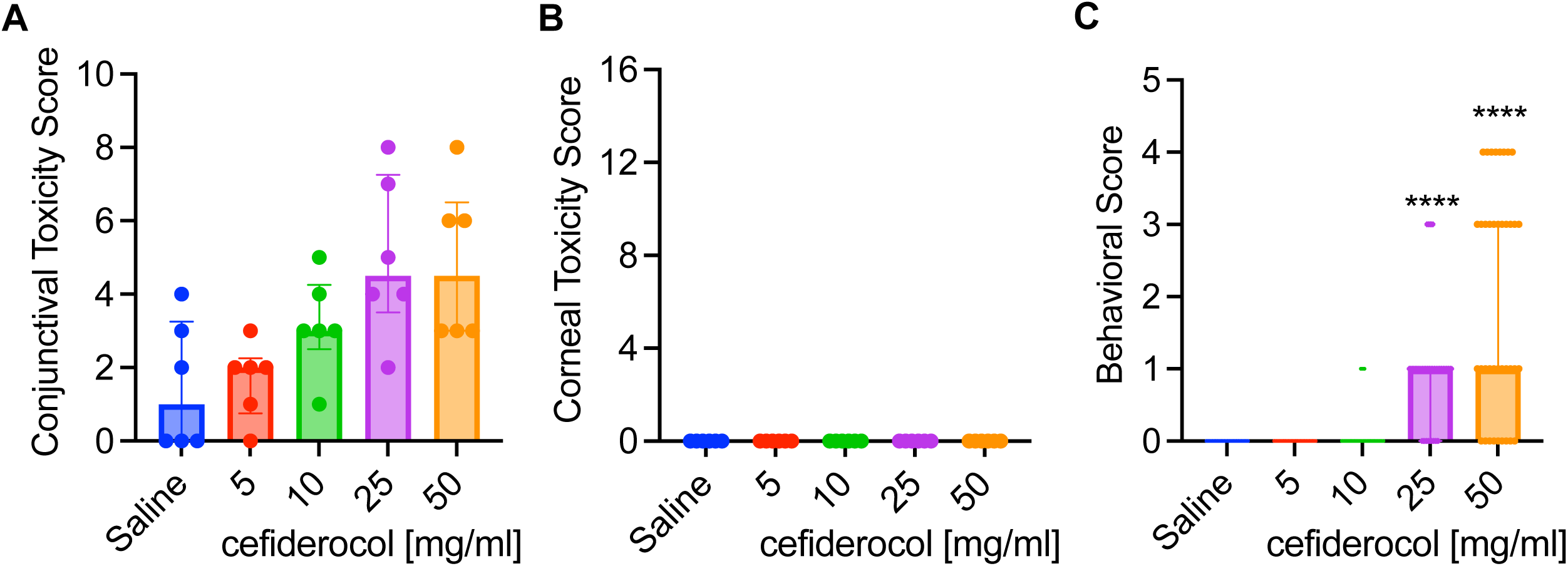
Tolerability and behavioral impact of cefiderocol treatment. **A-B.** Toxicity scores (max 10 for conjunctival, 16 for corneal) of NZWR exposed to topical drops of cefoderocol. Medians and interquartile ranges are shown. n=6. No statistical differences were found between groups. **C.** Behavioral signs of NZW rabbits exposed to cefoderocol topical drops. Medians and interquartile ranges are shown, n=39. Asteristiks inticate significant differences from saline group by Kruskal Wallis with Dunn’s Post-test, p<0.001.

Instillation of the 50 mg/ml and 25 mg/ml cefiderocol solutions produced more behavioral signs from the rabbits than the 10 mg/ml and 5 mg/ml cefiderocol solutions as well as the saline control over the course of the dosing. Based on the scoring system, both the 50 mg/ml and 25 mg/ml cefiderocol solutions produced a median score of 1.0 which equates to eye closing after instillation (Figure 3C). This is the minimal score after no reaction. Infrequently, there were other behavioral signs such as eye wiping and flinching demonstrated in the 50 mg/ml group as well as eye wiping in the 25 mg/ml group.

There was resolution of all conjunctival findings from day 0 to day 3 suggesting there was no delayed ocular toxicity associated with topical cefiderocol treatment. Since the maximum tested cefiderocol concentration was well tolerated and was same concentration as other cell wall targeting antibiotics (cefazolin, ceftazidime, and vancomycin), the highest concentration tested, 50 mg/ml, was selected for use in the keratitis studies.

### Efficacy of Cefiderocol against an Antibiotic-Susceptible *P. aeruginosa* Isolate in the NZW Rabbit Keratitis Model

The majority of *P. aeruginosa* keratitis isolates are not multidrug resistant ^10, 27^. Therefore, we first employed an antibiotic susceptible strain, K900 (Table 1) in our NZW rabbit keratitis model. These corneas were injected with a mean of 2350 ± 1414 CFU of *P. aeruginosa* K900 for the two trials. At 16 h post-infection, CFU from one group was measured prior to treatment (Onset). After 8 hours of topical treatment CFU were evaluated (Figure 4A). There was no increase in CFU in the saline group compared to the Onset for this strain suggesting that the maximum bacterial burden had been achieved. All antibiotic treatment groups had significantly reduced CFU compared to the saline control group. These decreases reached the level of bactericidal (99.9%) reductions compared to the Onset control. Furthermore, 67% of the corneas were sterilized with cefiderocol and ciprofloxacin treatments, while 50% of the corneas were sterilized with tobramycin treatment. There were no significant differences in CFU, or the number of eyes sterilized between the three antibiotic treatments.

**Figure 4.**
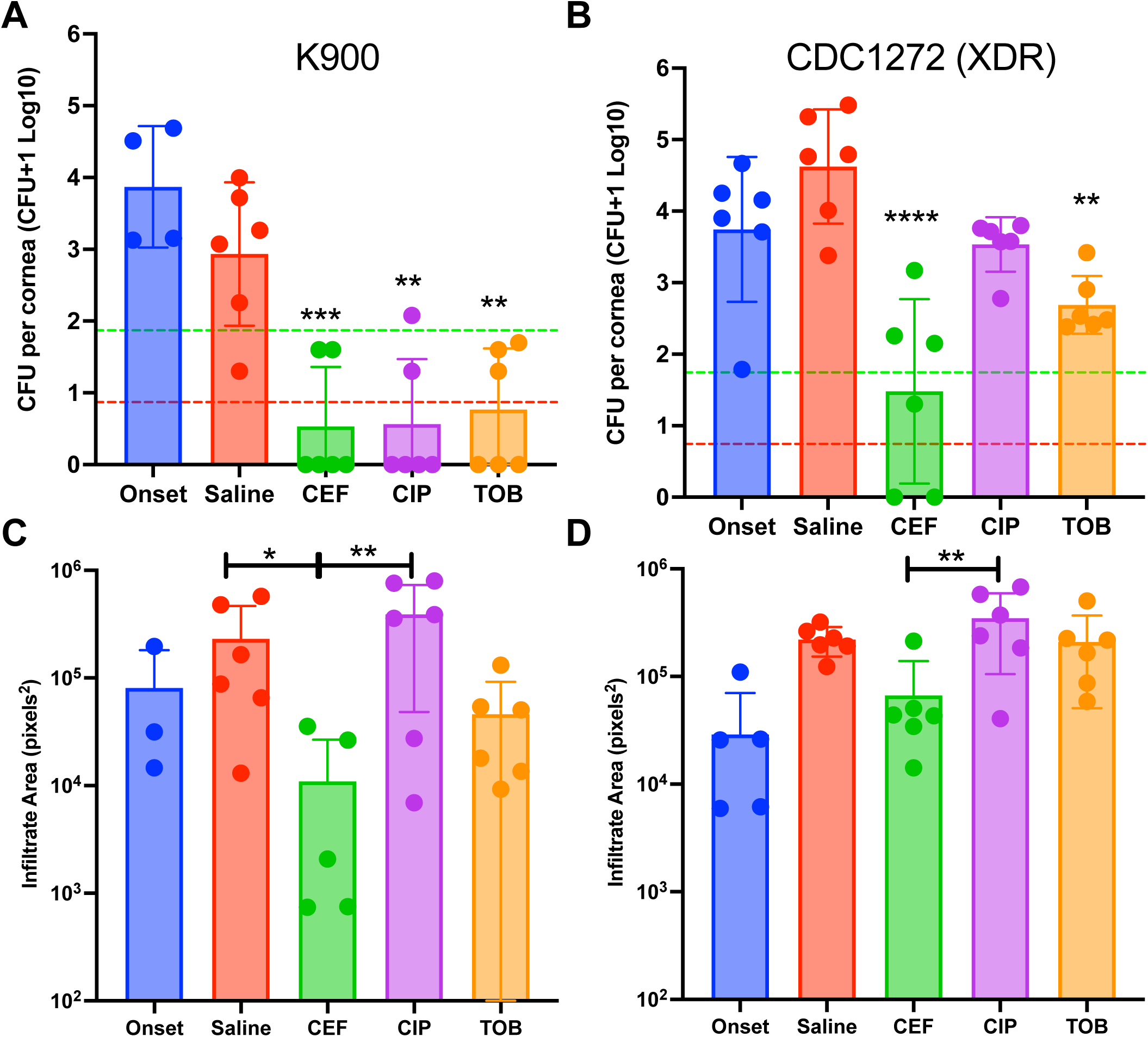
Efficacy of antibacterial treatment. **A-B.** The CFU per cornea were determined prior to treatment (Onset) or after 8h of topical treatment. Asterisks indicated significant differences from the saline group by ANOVA with Tukey’s Post-test. The pistacio lines represent a 99%, and the red lines a 99.9% reducion in CFU compared to the onset. n=6 except for onset group n≥3. **C-D.** Images of eyes were taken prior to euthanasia from a fixed distance.The onset group was imaged 16 h post infection, and before treatment. The other groups had 16 h of untreated infection, followed by 8 hours of treatment as indicated. Relative infiltrate area was determined using ImageJ software. Mean and SD are shown and were compared by ANOVA and Tukey’s post-test. *, p<0.05; **, p<0.01; ***, p<0.001; ****, p<0.0001.

### Antibacterial efficacy of Cefiderocol against an Extensively-Drug Resistant *P. aeruginosa* Keratitis Isolate in the NZW Rabbit Keratitis Model

By contrast to strain K900, the CDC1270 strain from the artificial tears outbreak was an XDR strain (Table 1). CDC1270 was used in the NZW rabbit keratitis model as was K900 above. A mean of 2325 ± 955 CFU of CDC1270 were injected for the two trials. The CFU at the Onset of therapy was nearly identical between strains. Unlike strain K900, the CDC1270 strain continued to proliferate by approximately 10-fold over the 8 hour treatment period in the saline group. Both cefiderocol and tobramycin significantly reduced CFU (Figure 4B) compared to the saline control whereas ciprofloxacin did not significantly reduce CFU. Cefiderocol was also significantly more effective than both ciprofloxacin and tobramycin in reducing corneal colony counts of XDR *P. aeruginosa*. Cefiderocol achieved a bactericidal (>99.9%) reduction compared to the saline treatment group but not the Onset group (>99%). Tobramycin did not achieve a 99% decrease in CFU compared to the saline control group. Moreover, cefiderocol sterilized 33% of the corneas compared to 0% for tobramycin and ciprofloxacin.

### Corneal Infiltrate Size

The areas of corneal infiltrates were measured from photographs taken from each eye prior to euthanasia (Figures 4C and 4D). In both K900 and CDC1270, infected eyes treated with ciprofloxacin trended toward being larger than the saline control and smaller than saline when treated with tobramycin and cefiderocol (Figures 4C and 4D). In all cases the cefiderocol treated eyes had significantly smaller infiltrates compared to the ciprofloxacin treated eyes (p<0.05).

### Approximate Corneal Cefiderocol Concentrations

The mean and standard deviation cefiderocol concentration obtained from the corneal homogenates from eyes infected with K900 and CDC1270 was 23.88 ± 8.69 µg/ml. The concentrations ranged from 12.53 µg/ml to 38.58 µg/ml per cornea. The mean cefiderocol corneal concentration was 191x greater than the MIC_90_ (0.125 µg/ml) and 25x greater than the MIC_100_ (0.94 µg/ml) of the *P. aeruginosa* keratitis isolates tested. Furthermore, the entire range of cefiderocol corneal concentrations was well above the MIC_90_ and MIC_100_ values.

### Stability of Refrigerated 50 mg/ml Cefiderocol

The results of stability testing of refrigerated 50 mg/ml cefiderocol solutions are presented in Figure 5. The zone sizes for all cefiderocol concentrations and refrigeration times are essentially the same suggesting that there is no loss in bioactivity of refrigerated 50 mg/ml cefiderocol solutions for up to 33 days.

**Figure 5.**
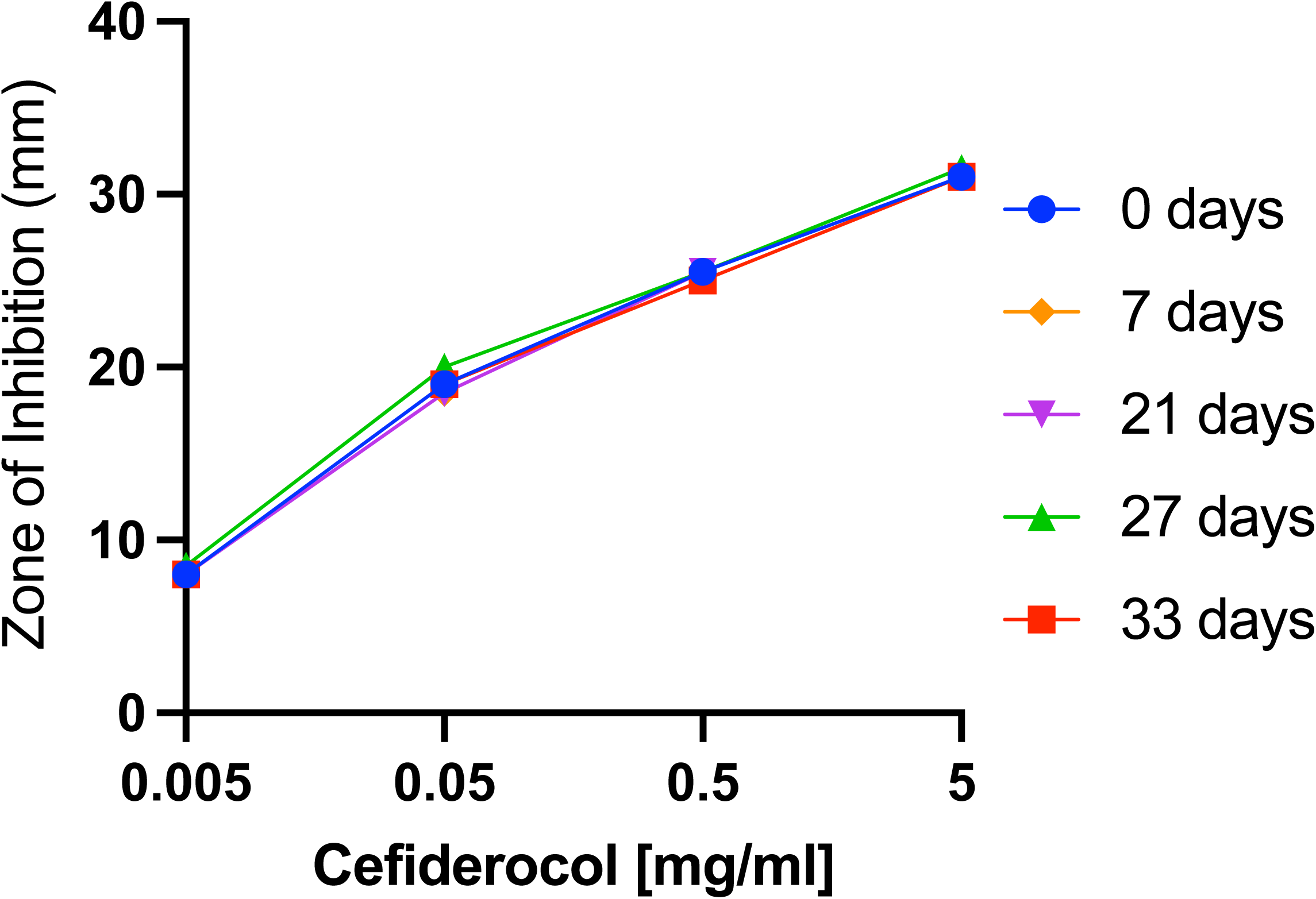
In vitro antimicrobial efficacy of solubilized cefiderocol stored at 4°C. Zones of inhibition of *P. aeruginosa* strain K900 are shown. Mean and SD of replicate experiments are shown.

## DISCUSSION

The reports of the recent outbreak of XDR *P. aeruginosa* ocular infections demonstrated a concerning lack of efficacy of conventional topical antibiotics. Several reports describe utilization of off label medications to attempt to control these aggressive infections. In two reports topically applied colistin by itself or in combination with fortified tobramycin-soaked collagen shields was used and resolved the infections after close to two months of therapy ^28, 29^. In another paper, a patient was treated topically with polymyxin B/trimethoprim and imipenem-cilastatin, but the infection was not resolved during the 1 month course of the published study ^3^. In another report, Morelli and colleagues ^2^, used intravenous cefiderocol in combination with topical application of imipenem-cilastatin and polymyxin B/trimethoprim and oral doxycycline. The patient’s infection appeared to resolve suggesting successful antibiotic treatment. As these reports demonstrate, there is no current consensus as to the most effective antimicrobial strategy to deal with XSR keratitis. Our current study was designed to test cefiderocol as a potential topical therapy for keratitis caused by XDR *P. aeruginosa*.

We observed that strains from our collection of keratitis isolates were more consistently susceptible to cefiderocol than ceftazidime, ciprofloxacin, or tobramycin. Notably the CLSI susceptibility breakpoints for tobramycin have recently been changed from 4.0 µg/ml to 1.0 µg/ml ^30^ which accounts for the high level of *in vitro* resistance recorded in this study. The systemic susceptibility breakpoints are not particularly useful for topical application because much higher local concentrations can be achieved in ocular tissue ^26, 31, 32^. However, strains such as the artificial tears outbreak XDR isolates have very high MIC values to commonly used antibiotics that may render them unable to reduce corneal bacteria even with topical drops. This study employed Mueller-Hinton II agar which is commonly used for MIC testing rather than iron-depleted media that can be used for testing susceptibility to cefiderocol ^33, 34^. This means that the measured values reported here may overestimate the MIC values.

50 mg/ml and 25 mg/ml cefiderocol were associated with conjunctival congestion, chemosis, and discharge; however, significant changes in the conjunctival signs were not observed. Most importantly, no corneal toxicity or fluorescein staining was produced for any cefiderocol concentration. The most common behavioral sign noted in the 50 mg/ml and 25 mg/ml concentrations was the closing of eyes after instillation. Occasionally, the rabbits wiped their eyes or even flinched after instillation in those groups. While these signs were not seen with the lower cefiderocol concentrations or the saline control they are similar to what has been previously reported with fortified antibiotics such as vancomycin. The pH of 50 mg/ml cefiderocol solution was acidic at 5.36, but is higher than other topical formulations such as ciprofloxacin 0.3% ophthalmic solution (pH 4.5) ^35^ and fortified vancomycin 50 mg/ml (pH ∼3.0) ^36^. There was no ocular toxicity or tolerability issues with the 10 mg/ml and 5 mg/ml cefiderocol concentrations. The data suggest that the tested formulations were well tolerated, but a specific ocular formulation with a more neutral pH may be advantageous. In this study however, we followed the manufacturer’s instructions for reconstitution and did not attempt to use a buffered solution for reconstitution to avoid any potential adverse effects on the activity or stability of cefiderocol. Based on the overall data from the ocular toxicity/tolerability study, we chose to use the highest cefiderocol concentration (50 mg/ml) for use in the NZW rabbit bacterial keratitis studies.

We employed a rabbit bacterial keratitis model that has previously been used for pre-clinical testing of antibiotics such as besifloxacin and gatifloxacin prior to their ophthalmic adoption. The XDR outbreak strain and a more typical antibiotic-susceptible *P. aeruginosa* keratitis isolate were used, as were standard of care antibiotics ciprofloxacin 0.3% and fortified tobramycin 14 mg/ml. Cefiderocol 50 mg/ml, ciprofloxacin 0.3%, and tobramycin 14 mg/ml showed equivalent efficacy against the antibiotic-susceptible strain K900. This is a similar outcome to a prior report for strain K900 and ciprofloxacin ^20^, and served to demonstrate that the ciprofloxacin and tobramycin antibiotics and treatment protocol used in this study were effective against a susceptible strain.

While all the antibiotic treatments indicated antibacterial efficacy against the antibiotic-susceptible *P. aeruginosa* strain K900, that was not the case with the XDR *P. aeruginosa* strain CDC1270. Cefiderocol produced a significant >3 Log_10_ decrease in colony counts compared to the saline control, a >2 Log_10_ decrease compared to ciprofloxacin and a > 1.4 Log_10_ compared to tobramycin. These differences were all significant. In addition, tobramycin 14 mg/ml also significantly decreased CDC1270 colony counts compared to the saline control.

Using a semi-quantitative method for determining cefiderocol concentrations, we determined that with aggressive topical treatment, the corneas achieved mean cefiderocol concentrations 25X higher than the MIC_100_ of the *P. aeruginosa* keratitis isolates tested in this study. These results demonstrate that high concentrations of cefiderocol can be achieved in corneas in which the corneal epithelium has been largely removed, which is the usual case with patients presenting with *P. aeruginosa* corneal ulcers.

We generally find no differences between the clinical signs produced among the various treatments in *P. aeruginosa* keratitis studies. This is due to clinical signs of infection being already present at the Onset of Therapy, and the short treatment duration (8 hours). However, during a review of the photographs of the bacterial keratitis studies, one notable clinical finding was observed. Corneal infiltrate areas were smallest in the cefiderocol treated eyes in all experiments suggesting that it is not antagonistic to healing or pro-inflammatory.

When designing this study, we had a concern about the stability of the cefiderocol solution. The product insert for FETROJA states that reconstituted solution can be stored at room temperature for up to 1 hour in the vial. Furthermore, it is recommended that FETROJA solution in the vial be immediately diluted. Diluted FETROJA was said to be stable for up to 6 hours at room temperature, and up to 24 hours when refrigerated and protected from light. (FETROJA Product Insert FET-PI-03). Because of these statements, we immediately diluted the FETROJA from the vial to produce the 50 mg/ml cefiderocol solution. This solution was made on the days of the experiments, kept on ice and in foil covered tubes during the dosing, and all excess was stored in foil covered tubes in a refrigerator. We chose to assess the stability of the 50 mg/ml cefiderocol solutions using a bioassay to determine activity. We found that there was no loss in antibiotic activity up to 33 days when stored refrigerated and out of light. This is important because patients will be using the drops for several days. Typically, fortified antibiotics have an expiration of 72 hours after preparation. Additional studies investigating the stability of the cefiderocol solutions at other temperature and storage conditions are indicated to mimic the potential storage conditions in the patient setting.

Limitations of the study include only testing *P. aeruginosa*, whereas keratitis can be caused by a number of bacterial pathogens and our data cannot speak to cefiderocol’s activity against them. One clear limitation of cefiderocol is its lack of documented efficacy against Gram-positive bacteria such as *Staphylococcus aureus* which is a frequent cause of keratitis. A longer-term study showing final outcomes would also be helpful. Nevertheless, this study demonstrates that topical cefiderocol is effective for treatment of *P. aeruginosa* in an animal model that is predictive of patient efficacy. This was true for a non-antibiotic resistant strain and for an XDR *P. aeruginosa* isolate from the recent outbreak. In conclusion, this is the first study to evaluate the ocular toxicity and antibacterial efficacy of topical ocular cefiderocol. Our results suggest that further development of cefiderocol for use as a topical ophthalmic drop for treatment of XDR *P. aeruginosa* keratitis is warranted.

## Abbreviations and Acronyms

CDC: Center for Disease Control and Prevention
MIC: Minimum Inhibitory Concentration
NZW: New Zealand White
NIH: National Institutes of Health
PBS: Phosphate Buffered Saline
XDR: Extensively Drug Resistant
CLSI: Clinical and Laboratory Standards Institute

## Acknowledgements

This study was funded by the Charles T. Campbell Ophthalmic Microbiology Laboratory, National Institutes of Health grants R01EY027331 (to R.S.), and CORE Grant P30 EY08098 to the Department of Ophthalmology. The Eye and Ear Foundation of Pittsburgh and from an unrestricted grant from Research to Prevent Blindness, New York, NY provided additional departmental funding.

