## supplemental figure for "Cefiderocol is an effective topical monotherapy for experimental extensively-drug resistant *Pseudomonas aeruginosa* keratitis"

**Figure S1.** This figure presents the eyes from each treatment group of the Cefiderocol Ocular Toxicity/Tolerability Study on the day of treatment. Subject numbers and right or left eyes are indicated above the photographs. The fluorescein stains eyes are below the normal photographs for each eye.

50 mg/ml Cefiderocol

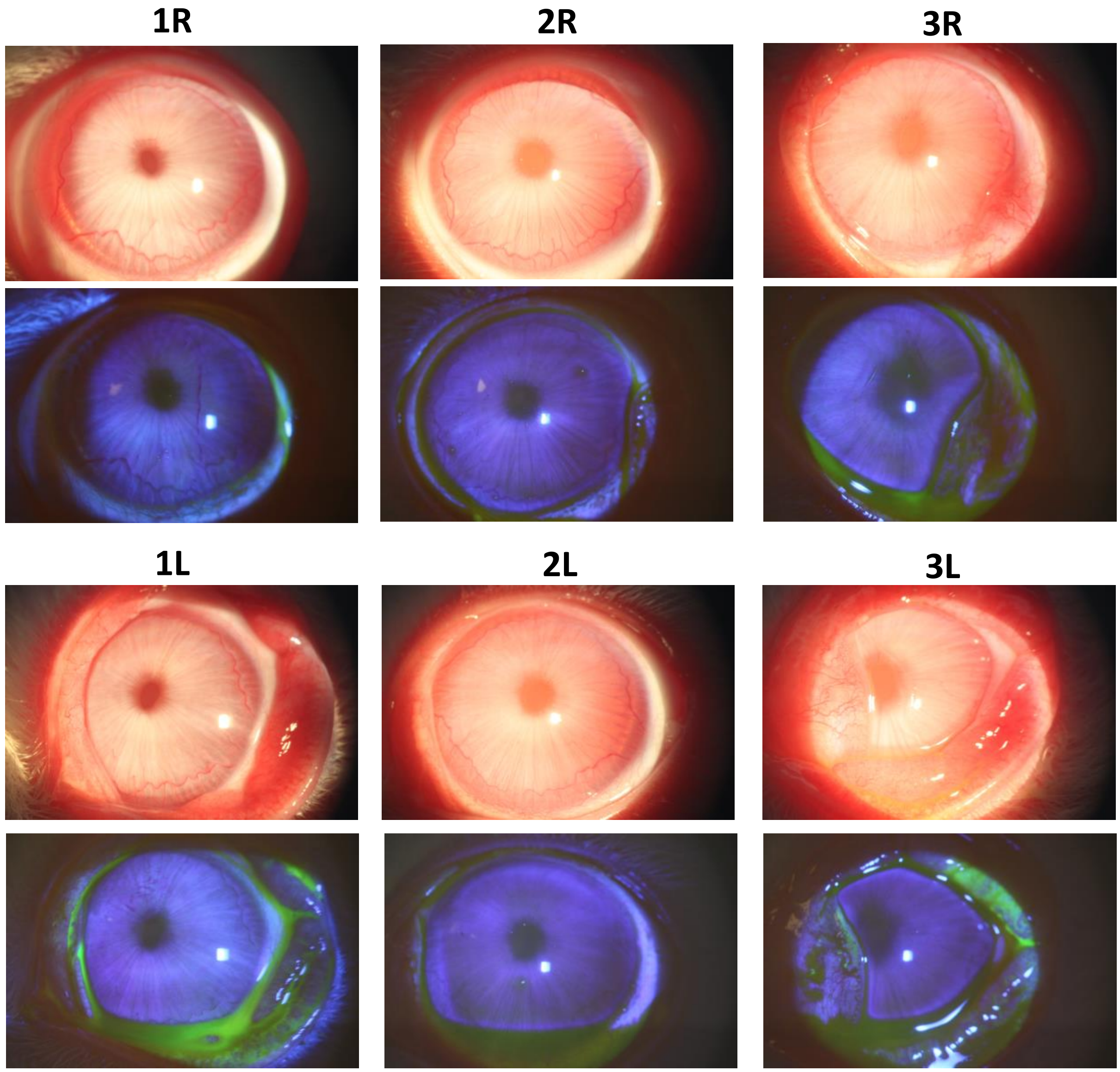

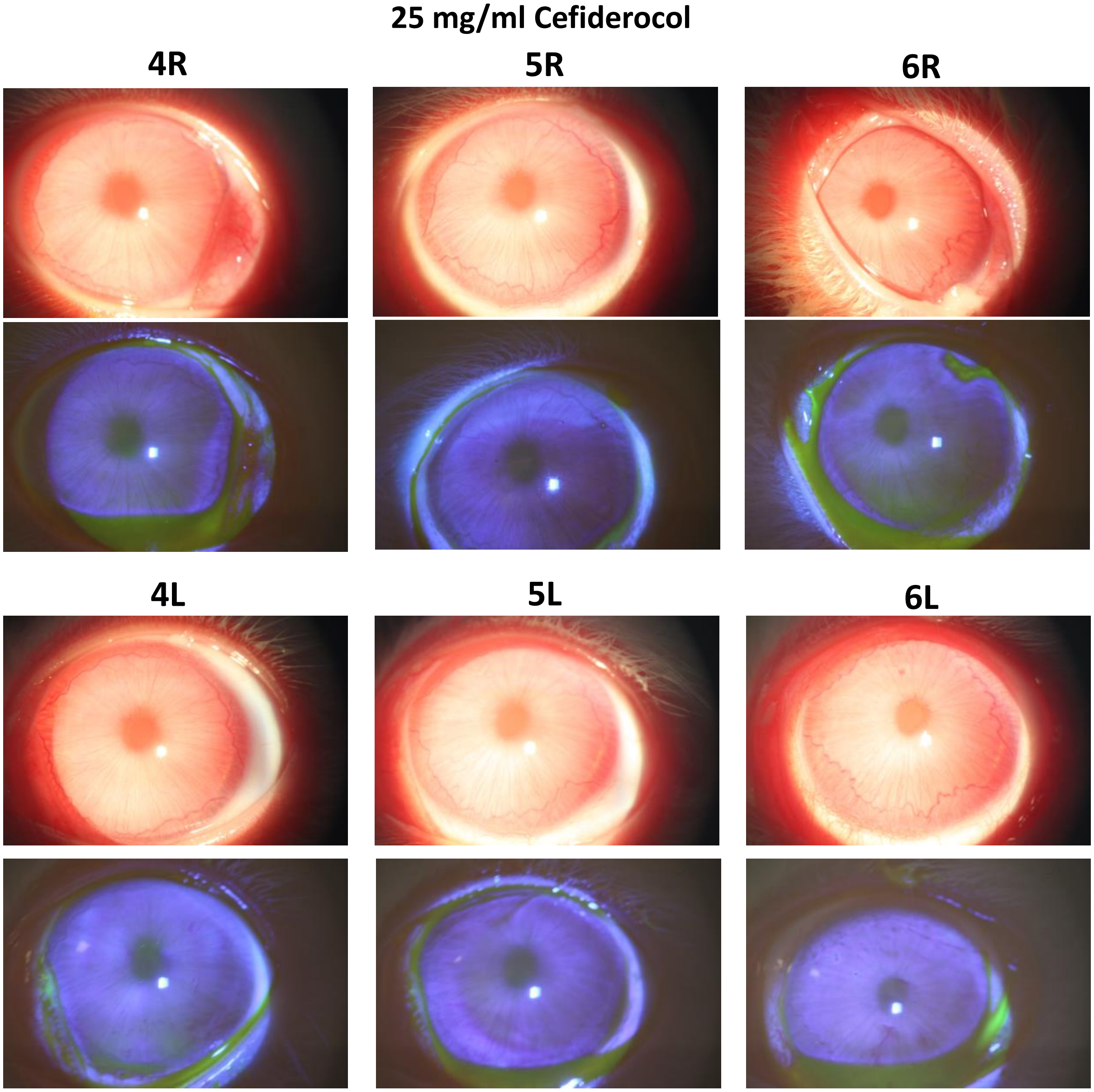
25 mg/ml Cefiderocol
4R
5R
6R
4L
5L
6L

10 mg/ml Cefiderocol

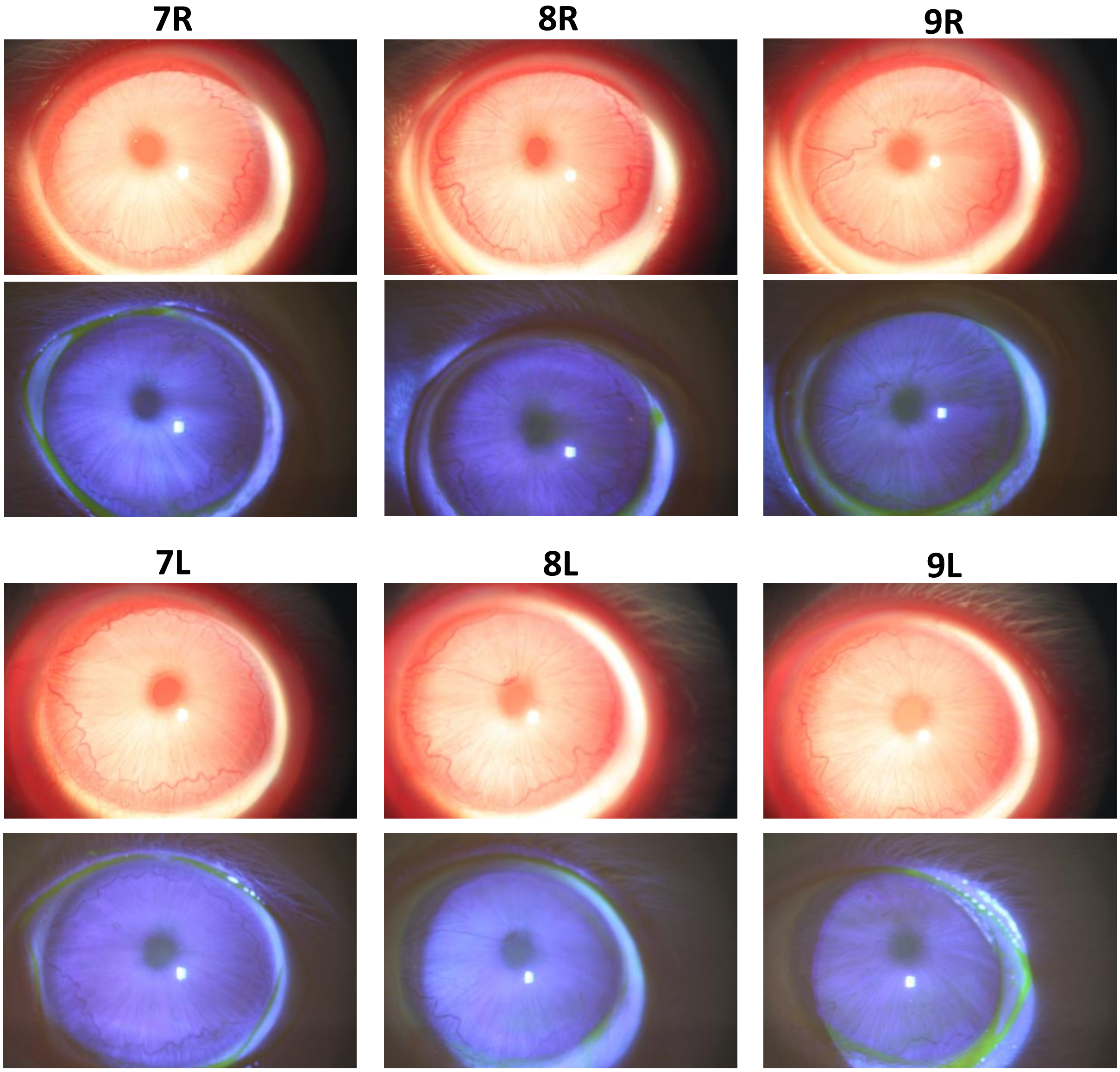

5 mg/ml Cefiderocol

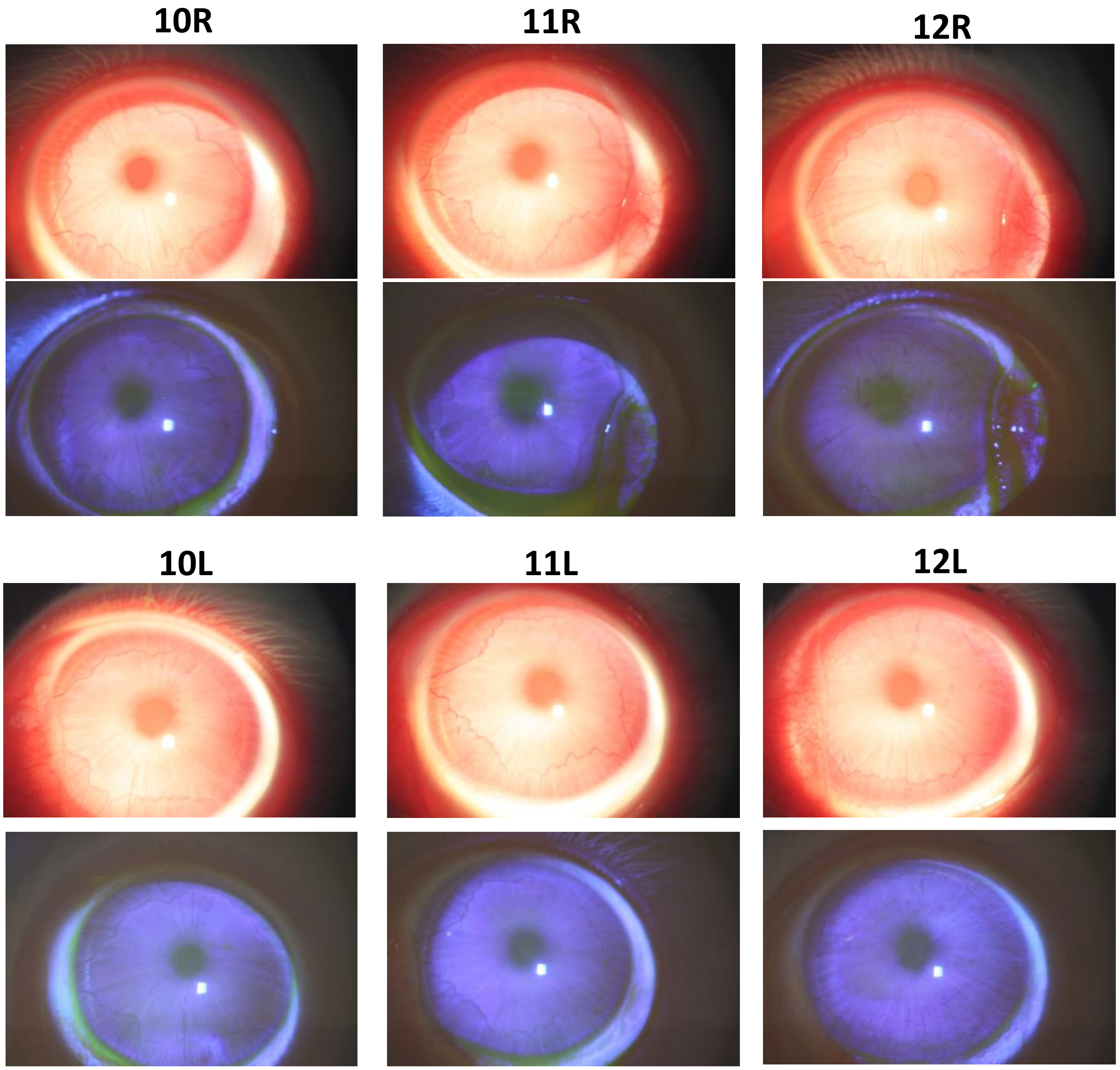

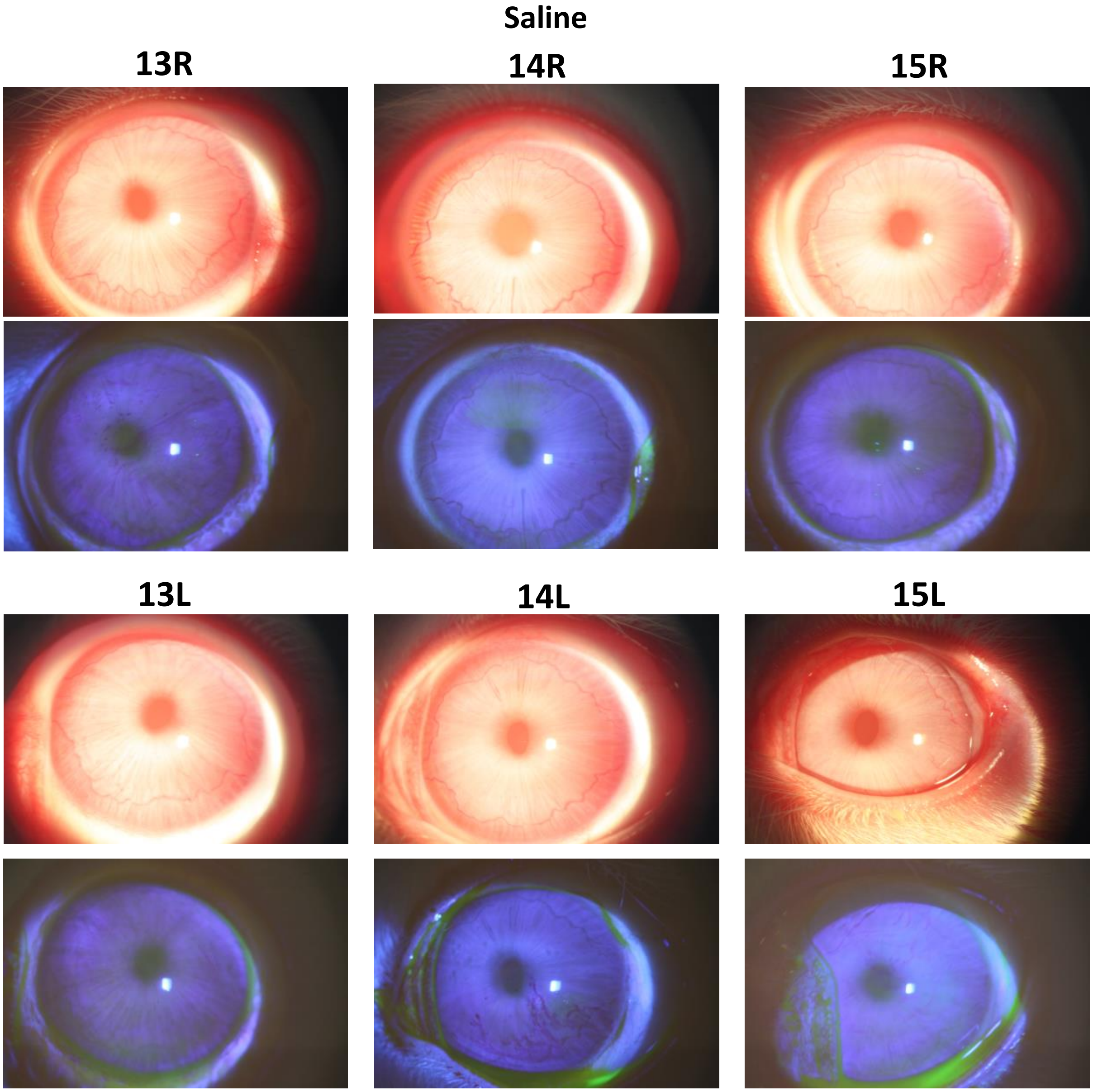
Saline
13R
14R
15R
13L
14L
15L
